# Full-length transcript characterization of *SF3B1* mutation in chronic lymphocytic leukemia reveals downregulation of retained introns

**DOI:** 10.1101/410183

**Authors:** Alison D. Tang, Cameron M. Soulette, Marijke J van Baren, Kevyn Hart, Eva Hrabeta-Robinson, Catherine J. Wu, Angela N. Brooks

**Affiliations:** Department of Biomolecular Engineering, University of California, Santa Cruz, CA; Department of Molecular Cell & Developmental Biology, University of California, Santa Cruz, CA; Department of Medical Oncology, Dana-Farber Cancer Institute, Boston, MA

**Author notes:** Correspondence: Angela N. Brooks.

## Abstract

*SF3B1* is one of the most frequently mutated genes in chronic lymphocytic leukemia (CLL) and is associated with poor patient prognosis. While alternative splicing patterns caused by mutations in *SF3B1* have been identified with short-read RNA sequencing, a critical barrier in understanding the functional consequences of these splicing changes is that we lack the full transcript context in which these changes are occurring. Using nanopore sequencing technology, we have resequenced full-length cDNA from CLL samples with and without the hotspot *SF3B1* K700E mutation, and a normal B cell. We have developed a workflow called FLAIR (Full-Length Alternative Isoform analysis of RNA), leveraging the full-length transcript sequencing data that nanopore affords. We report results from nanopore sequencing that are concordant with known *SF3B1* biology from short read sequencing as well as altered intron retention events more confidently observed using long reads. Splicing analysis of nanopore reads between the *SF3B1*^*WT*^ and *SF3B1*^*K700E*^ samples identifies alternative upstream 3’ splice sites associated with *SF3B1*^*K700E*^. We also find downregulation of intron retention events in *SF3B1*^*K700E*^ relative to *SF3B1*^*WT*^ and no difference between CLL *SF3B1*^*MT*^ and B cell, suggesting an aberrant intron retention landscape in CLL samples lacking *SF3B1* mutation. With full-length isoforms, we are able to better estimate the abundance of RNA transcripts that are productive and will likely be translated versus those that are unproductive. Validation from short-read data also reveals a strong branch point sequence in these downregulated intron retention events, consistent with previously reported branch points associated with mutated *SF3B1*. As nanopore sequencing has yet to become a routine tool for characterization of the transcriptome, our work demonstrates the potential utility of nanopore sequencing for cancer and splicing research.

---

Mutations in the splicing factor *SF3B1* in various cancers have been associated with characteristic alterations in splicing. Specifically, recurrent somatic mutations in *SF3B1* have been linked to various diseases including chronic lymphocytic leukemia (CLL) and myelodysplastic syndromes, with HEAT-repeat domain mutations being implicated in contributing to poor clinical outcome [1–4]. SF3B1 is a core component of the spliceosome and associates with the branch point (BP) adenosine, stabilizing the interaction between the U2 snRNA/BP complex [5–7]. Mutations in *SF3B1* induce aberrant splicing patterns that have been well-characterized using short-read sequencing (∼100nt) of the transcriptome. In particular, the K700E hotspot mutation in the HEAT-repeat domain of *SF3B1* has been associated with altered splice sites at the 3’ end of introns [8–10]. SF3B1^K700E^ affects acceptor splice sites due to changes in BP recognition and usage [11–13].

While alternative splicing patterns on a junction-level have been observed, these patterns have not been systematically examined on an isoform level, which limits our complete understanding of the functional consequences of these aberrant splicing changes. Due to limitations in short-read sequencing technology, an incomplete picture of the transcriptome is assembled because exon connectivity is difficult to determine from fragments [14]. Moreover, detection and quantification of transcripts containing retained introns is difficult and often dismissed [15]. However, isoform-level analyses are imperative for building a better understanding of altered transcripts in a cell. Long-read sequencing techniques offer increased information on exon connectivity by sequencing full-length transcripts [16–18]. In particular, nanopore minION sequencing yields long reads up to 2 megabases [19], which has been used for applications such as sequencing the centromere of the Y chromosome [20], the human genome [21], and single-cell transcriptome sequencing [22]. By converting changes in current caused by blockage of DNA threading through a nanopore into sequence, nanopore sequencing can sequence complete molecules of DNA [23]. Nanopore technology has yet to be thoroughly explored as a tool for investigating changes in splicing in the context of cancer-associated splicing factor mutations. Using nanopore sequencing technology, we have resequenced two chronic lymphocytic leukemia (CLL) patient tumor samples and one normal B cell. We use nanopore reads to investigate alternative splicing events associated with *SF3B1* K700E mutation. We use a hybrid-seq approach, combining the accuracy of Illumina short reads for splice junction accuracy with the scaffolding power of long reads to overcome the higher error rates of long reads. Previous PacBio hybrid-seq approaches rely more heavily on short reads or disregard raw reads with less than 99% accuracy [24,25]. Our workflow, FLAIR (Full-Length Alternative Isoform analysis of RNA), assembles complete isoforms from higher-error nanopore reads with high-accuracy at splice junctions for consequent analysis of alternative isoform usage between sequenced samples. Splicing analysis of nanopore reads identifies a bias toward increased alternative 3’ splice sites (3’SS) than alternative 5’ splice sites (5’SS) in the *SF3B1*^*K700E*^ sample. We also highlight a previously underappreciated finding of differential intron retention in *SF3B1*^*K700E*^ versus *SF3B1*^*WT*^ CLL that suggests an aberrant intron retention landscape in wildtype *SF3B1* samples and increased splicing in *SF3B1*^*K700E*^ samples.

## Results

### Full-length alternative isoform analysis of RNA from chronic lymphocytic leukemia

To characterize *SF3B1*^*K700E*^ full-length transcripts, we performed nanopore sequencing in a primary CLL *SF3B1*^*K700E*^ sample that was clonal for the mutation and a CLL *SF3B1*^*WT*^ sample that did not contain other common CLL-associated mutations. Both of these samples were previously analyzed in a study to characterize splicing changes associated with *SF3B1* mutation, using short-read RNA-Seq data [8]. We obtained 119,091, 120,798, and 46,077 reads using nanopore 2D sequencing technology for the CLL *SF3B1*^*WT*^ sample, CLL *SF3B1*^*K700E*^ sample, and a B cell sample, respectively (Table 1). Many existing tools for short-read RNA-Seq analysis lack the ability to process long reads or reads with high error profiles. As such, we developed a workflow to analyze long nanopore reads for splicing changes called FLAIR (Full-Length Alternative Isoform analysis of RNA) (Fig. 1). First, we align the raw read sequences from all samples to the genome to identify the general transcript structure (Fig. 1A). To evaluate existing spliced alignment tools with nanopore sequence data, we compared the long-read spliced-aligner, minimap2 [26], against GMAP [27], which has been used in previously reported long-read studies [22,28,29]. The aligners were evaluated according to splice site accuracy, with minimap2 demonstrating marked improvement in splice site mapping when comparing aligned sites with annotated sites (GENCODE v24, Fig. 2A). As *SF3B1* mutations alter 3’ splice site choice [8–10,13], we increased splice site accuracy by correcting read alignments based on short-read evidence (Fig. 2B). The high frequency of deletions in nanopore reads causes many alignment gaps, which we resolve by smoothing gaps of 30 nt or fewer. High error rates and deletions in sequencing can cause spurious alignments near annotated splice sites or splice sites with support from short-read data (Fig. 1B, Supplementary Fig. 1A,B). We corrected reads with misaligned splice sites to the nearest splice site within a window size of 10 base pairs, using evidence from matched short-read data and annotated comprehensive GENCODE junctions (Supplementary Fig. 1C,D). 93.17% (687,980 of 738,442) of all sites were mapped perfectly, i.e. matching annotated and short-read supported splice sites, not needing correction. Sites that align outside of the wiggle room to canonical splice motifs with 3 supporting nanopore reads, but no annotation or short-read support, are deemed novel. After splice site correction, we identified 1,839 novel splice sites present in our nanopore reads. Upon further manual inspection, these novel sites appeared to be driven by alignment errors of shorter exonic regions. Given the lower confidence in these novel splice sites, we removed them from our further analyses.

**Table 1:** Nanopore sequencing statistics.

| Sample | 2D pass reads | % 2D aligned reads | Genes covered | Error rate |
| --- | --- | --- | --- | --- |
| B cell | 46,077 | 69.3 | 4,945 | 11.004% |
| CLL SF3B1<sup>WT</sup> | 119,091 | 60.6 | 7,085 | 14.512% |
| CLL SF3B1<sup>K700E</sup> | 120,798 | 75.9 | 8,803 | 12.605% |
| Simulated | 117,512 | 92.7 | 7,361 | 16.824% |

**Figure 1:**
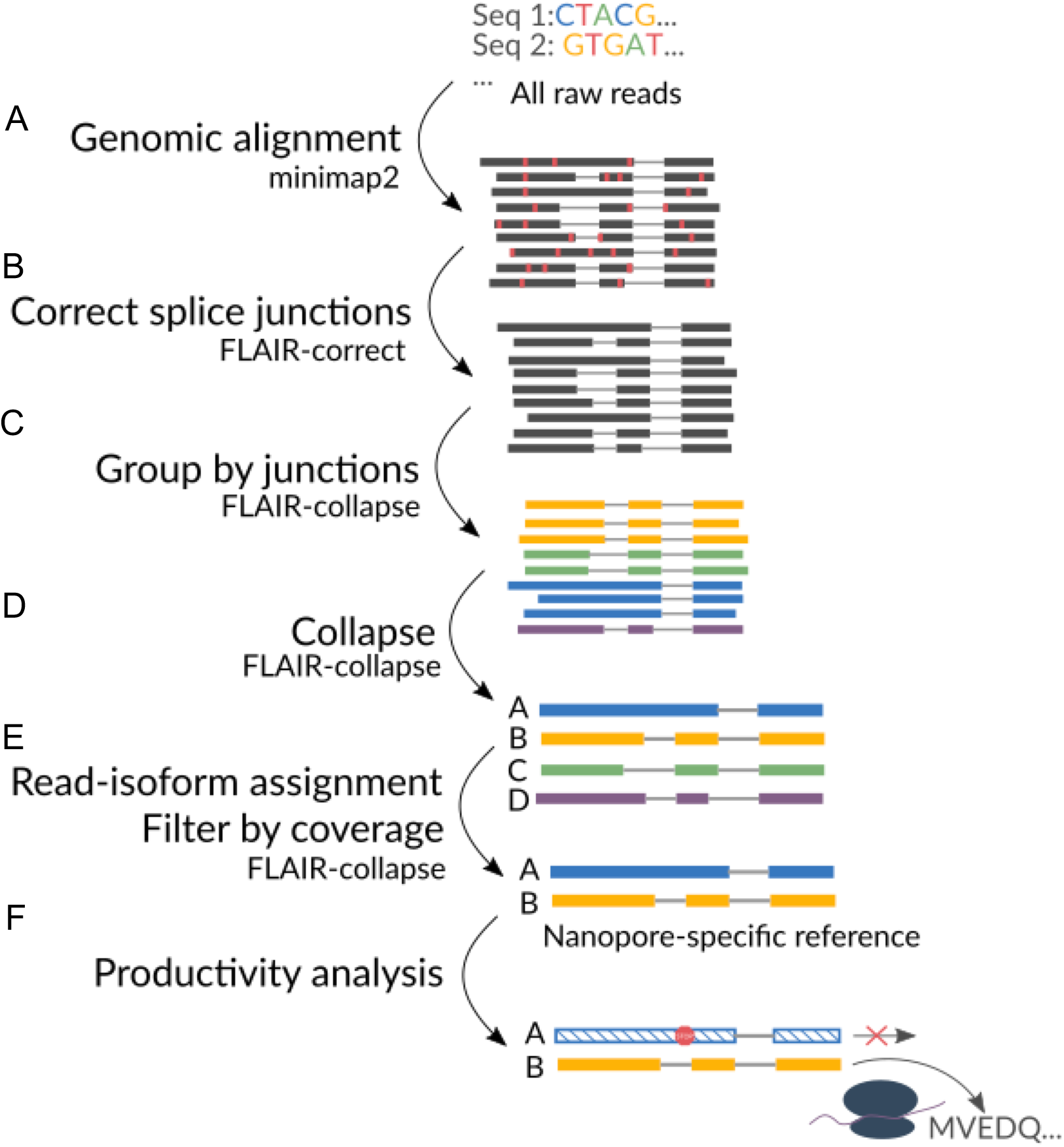
FLAIR workflow for nanopore isoform assembly and splicing analysis. A.Raw reads are aligned to the genome using minimap2. B.Read errors are corrected, and misaligned junctions are corrected using junctions observed in matched short-read and genomic annotation. C.Isoforms are grouped based on splice junctions (colors denote each group). D.Isoform groups are collapsed into a first-pass assembly of representative isoforms where the most commonly used transcription start and ends (within a 20 nt window) are used to represent the isoform. E.Raw reads are realigned to the first-pass assembly from D, and each representative isoform’s expression is quantified. Isoforms with supporting reads greater than a specified threshold are kept in the final set of isoforms, called the nanopore-specific reference. F.The nanopore-specific reference produced in 1E can be further quantified, examined for alternative splicing, and assessed for productivity.

**Figure 2:**
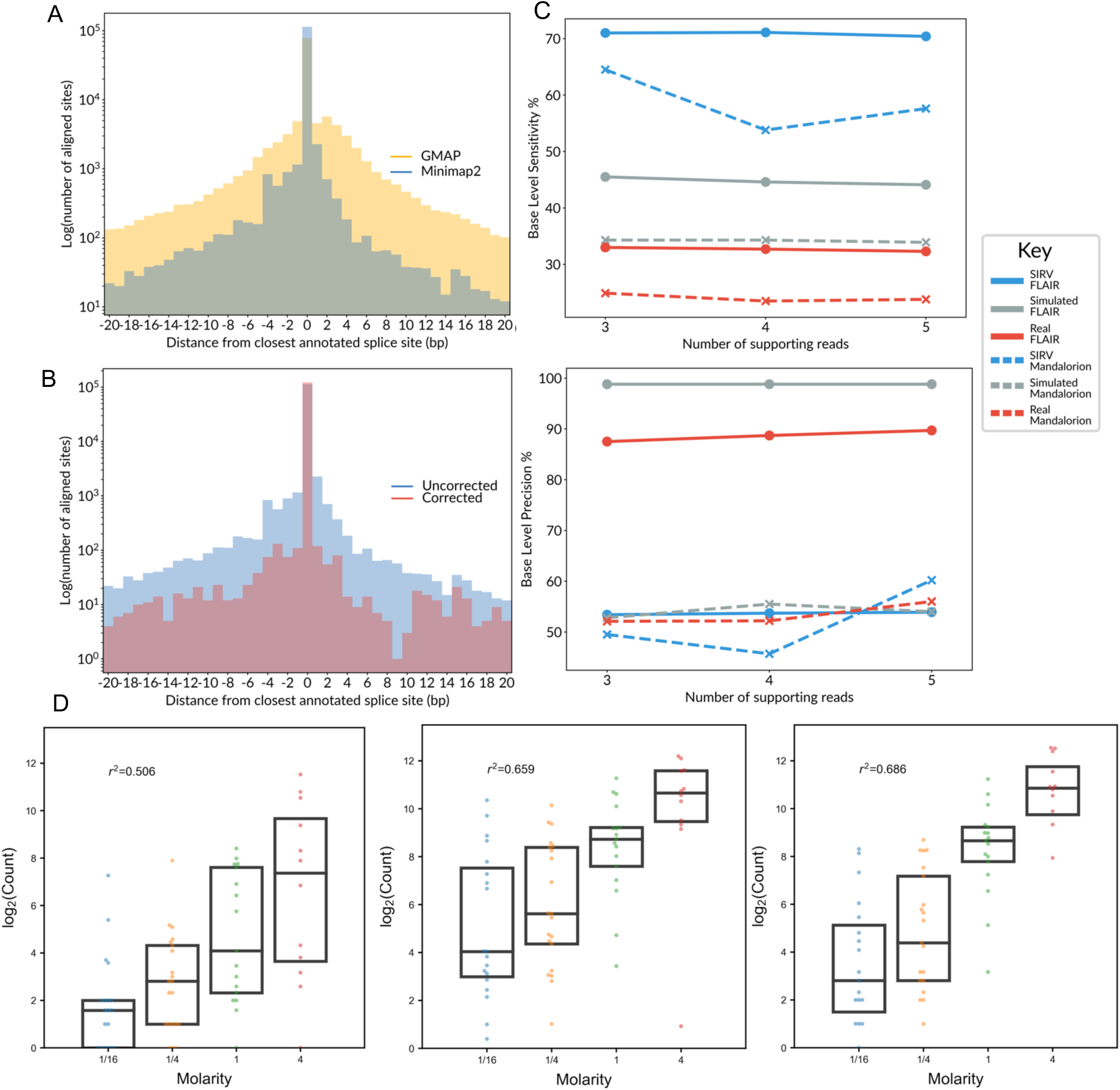
Evaluation of the FLAIR workflow. A.Bar plot showing splice site accuracy comparison for GMAP and minimap2 alignments. X-axis is the distance from the aligned site to the closest annotated site, the y-axis is the number of aligned sites normalized and log-scaled. B.Same as A, showing splice site accuracy comparison between raw and splice corrected minimap2 alignments. C.Base level precision (top) and sensitivity (bottom) for isoform assembly method comparison between Mandalorion (dashed) and FLAIR (solid). Datasets used are SIRV 2D (blue), simulated CLL data (orange), and real CLL data (blue. X-axis is the minimum number of supporting reads thresholds tested. D.Quantification-method evaluation comparing counts from uniquely aligned reads only, weighted identity, and primary read alignment counting (left to right) using SIRV data. R^2^ are Pearson correlation values.

To perform isoform-level analyses, we identified a set of expressed isoforms with high confidence. Initially, FLAIR creates a first-pass nanopore isoform assembly by grouping all reads with the same unique set of splice junctions used into isoform groups (Fig. 1C). To do this, we used splice-corrected nanopore reads to group isoforms by the splice junctions they contained. Within isoform groups, we called confident transcription start and end sites based on the density of read start and end positions. The first-pass isoform assembly produces a mixed set of isoforms found in annotations as well as isoforms with splicing patterns only observed in nanopore reads. We then quantified the number of reads aligning to each isoform by realignment of raw reads to the first-pass assembly. This realignment of reads to the set of nanopore-specific isoforms allowed us to account for misalignments that manifested during genomic alignment and more correctly assign reads to isoforms. The realignment was also crucial for better distinguishing splice site differences, as splice site differences caused by *SF3B1* mutation can cause subtle differences from canonical sites of only a few basepairs [8]. In the final step, we produced a confident nanopore isoform assembly by filtering for isoforms that passed a threshold with a minimum of 3 supporting reads (Supplementary File 1).

### FLAIR provides improved full-length isoform detection from long-reads with sequencing errors

We evaluated FLAIR against another nanopore isoform-building method, Mandalorion [22]. We benchmarked using gffcompare [30] on assemblies using both Lexogen Spike-In RNA Variants (SIRV) sequenced with nanopore 2D technology, a simulated dataset, and using our real nanopore dataset against GENCODE transcripts. SIRV transcripts are of known sequence and concentration and are of comparable complexity to human transcripts [31]. We simulated nanopore transcriptome reads by creating a wrapper for NanoSim [32], a tool used for simulating genomic nanopore reads (Methods). We found FLAIR to have increased sensitivity and precision at varying read depths using our three different data types and we observed nominal changes in sensitivity and precision at different read thresholds (Fig. 3C). As a result, we decided to use the lowest threshold of 3 to maximize the number of isoforms assembled. Using FLAIR, we identified a total of 10,613 high-confidence spliced isoforms. 40.7% of all reads were full-length, with full-length defined as spanning 80% of the high-confidence assembled isoforms. Although the RNA samples had been stored for three years, they showed minimal signs of degradation as this percentage of full-length reads is comparable to what is observed from other nanopore sequencing data [16].

**Figure 3:**
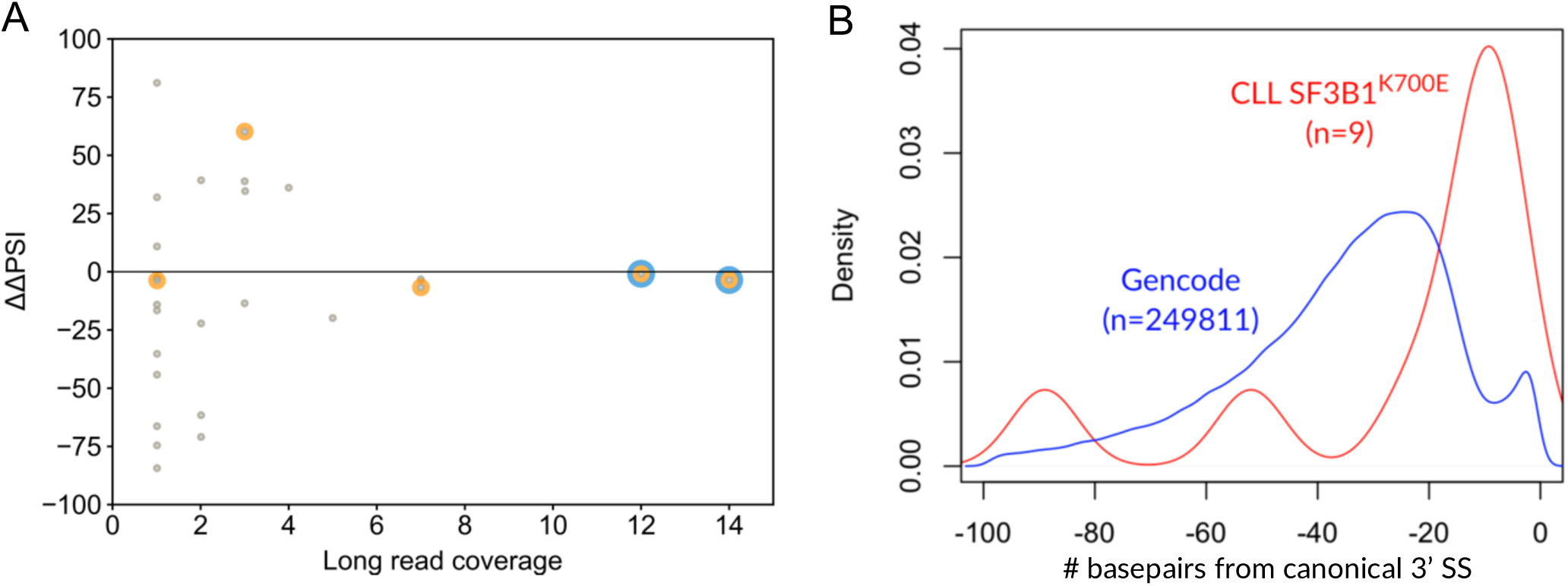
Alternative 3’ splice site usage associated with *SF3B1* mutation. A.Splicing event nanopore read coverage versus ΔΔPSI plot. Each point in grey is an altered 3’ splice site associated with *SF3B1* mutation, identified through short read data. The x-axis is the total nanopore read coverage of the altered junction from the SF3B1^WT^ and SF3B1^K^700^E^ samples. Points with orange and blue circles indicate a minimum total coverage of the (aberrant junction + canonical junction) of 3 and 5, respectively, in either the SF3B1^WT^ or SF3B1^K^700^E^ sample. The y-axis is the difference of ΔPSI values (SF3B1^K^700^E^ PSI - SF3B1^WT^ PSI) in the short versus long read data. B.Distance between the alternative 3’ SS between SF3B1^K^700^E^ and SF3B1^WT^ (p < 0.1, red line) and canonical splice sites. The distance between canonical GENCODE v24 basic annotated 3’ SS to the first non-GAG trimer (blue line).

### Nanopore read analysis of SF3B1^K700E^ associated alternative isoform usage is consistent with short-read data

Quantifying expression of isoforms using nanopore data may appear to be straightforward in that each read corresponds to a complete transcript present in the cell; however, ambiguity of read assignments to transcripts can occur due to sequence errors, genes with increased isoform complexity, and reads that are not full-length. We benchmarked different approaches to assigning reads to transcripts using nanopore 2D-sequenced SIRVs. We evaluated three different approaches of isoform quantification, which include considering only uniquely mapped reads, weighting alignments to transcripts with higher identity scores and proportion of transcript length, or only primary read alignments. While the latter two methods are comparable, we proceeded with the consideration of only primary alignments in consequent analyses as the method performed better for the quantification of 2D SIRVs (Fig. 2D).

Previous literature has shown that SF3B1^K700E^ promotes alternative 3’ splice site usage [8], a pattern we sought to validate in our data. To evaluate the ability for nanopore sequencing to quantify alternative splicing patterns, we compared percent spliced-in (PSI) values between short-read and long-read data from significantly altered 3’ splicing events associated *with SF3B1*^*K700E*^ (Fig. 3A). We found that PSIs are more similar across the two sequencing technologies with increasing read depth; however, many splice junctions had insufficient coverage in our nanopore data for adequate power to detect the same splice site usage. Next, we measured the effect of SF3B1^K700E^ on altering acceptor and donor sites by comparing SF3B1^WT^ (combining both B-Cell and CLL SF3B1^WT^) to CLL SF3B1^K700E^. Using the nanopore data, we identified 7 alternative 3’ splice sites and 4 alternative 5’ splice sites that were differentially spliced (Fisher’s exact test, p<0.05) between *SF3B1*^*WT*^ and CLL *SF3B1*^*K700E*^ based on stringent filters for total read depth. We looked into the distances between altered 3’ splice sites (with a less stringent cutoff of p<0.1) and canonical splice sites and observed a distribution that peaks around -17 bp. A comparison of GENCODE canonical 3’ splice sites distances to the first non-GAG trimer showed a control distribution significantly different from SF3B1^K700E^-altered sites (Wilcoxon p=3.80e-4) (Fig. 3B). These distributions were concordant with previous reports showing that alternative 3’ splice sites in *SF3B1*^*K700E*^ appear to be upstream of those of *SF3B1*^*WT*^ [8]. The full distribution both up- and downstream of the canonical splice site of shows that more alterations cause upstream 3’ SS (Supplementary Fig. 2). As another control, we looked at altered 3’ SS between B cell and CLL SF3B1^WT^ and found only one splicing event to be altered (p<0.05), which is in agreement with our expectation in SF3B1^WT^ samples. Due to lack of coverage (Fig. 3A), these differential splicing changes were not significant given multiple testing; however, trends in changes in splicing were consistent with prior CLL SF3B1-associated biology.

### Down-regulated intron retention in SF3B1-mutated CLL is associated with a strong branch point sequence

Intron retentions have been observed to differentiate tumors from matched normal tissue, with intron retentions prevalent across a variety of cancers [33,34]. However, it is difficult to characterize intron retention event usage confidently using short reads without reads spanning the complete intron, including exon-intron junctions [15]. With long reads, intron retention events are better identified and quantified, regardless of the length of the intron. Quantifying expression of isoforms with and without retained introns between CLL *SF3B1*^*K700E*^ and *SF3B1*^*WT*^ revealed that isoforms with intron retention were globally down-regulated in the mutant sample compared to CLL *SF3B1*^*WT*^ after normalizing for library size (Fig. 4A). Comparing the CLL samples with a normal B-cell, the *SF3B1*^*WT*^ sample showed a relative increase in retained introns while the *SF3B1*^*K700E*^ showed no difference in expression of retained introns (Mann-Whitney U p=1.95e-4 and p=0.223) (Supplementary Fig. 3A,B). Reanalysis of short-read data from these same samples confirmed the observed increase in expression of intron-retaining (IR) isoforms in CLL *SF3B1*^*WT*^ samples (Fig. 4B). We next identified introns that were differentially retained between *SF3B1*^*K700E*^ and *SF3B1*^*WT*^ using nanopore sequencing data and found 25 introns (Fisher’s exact test, p-value <0.05).

**Figure 4:**
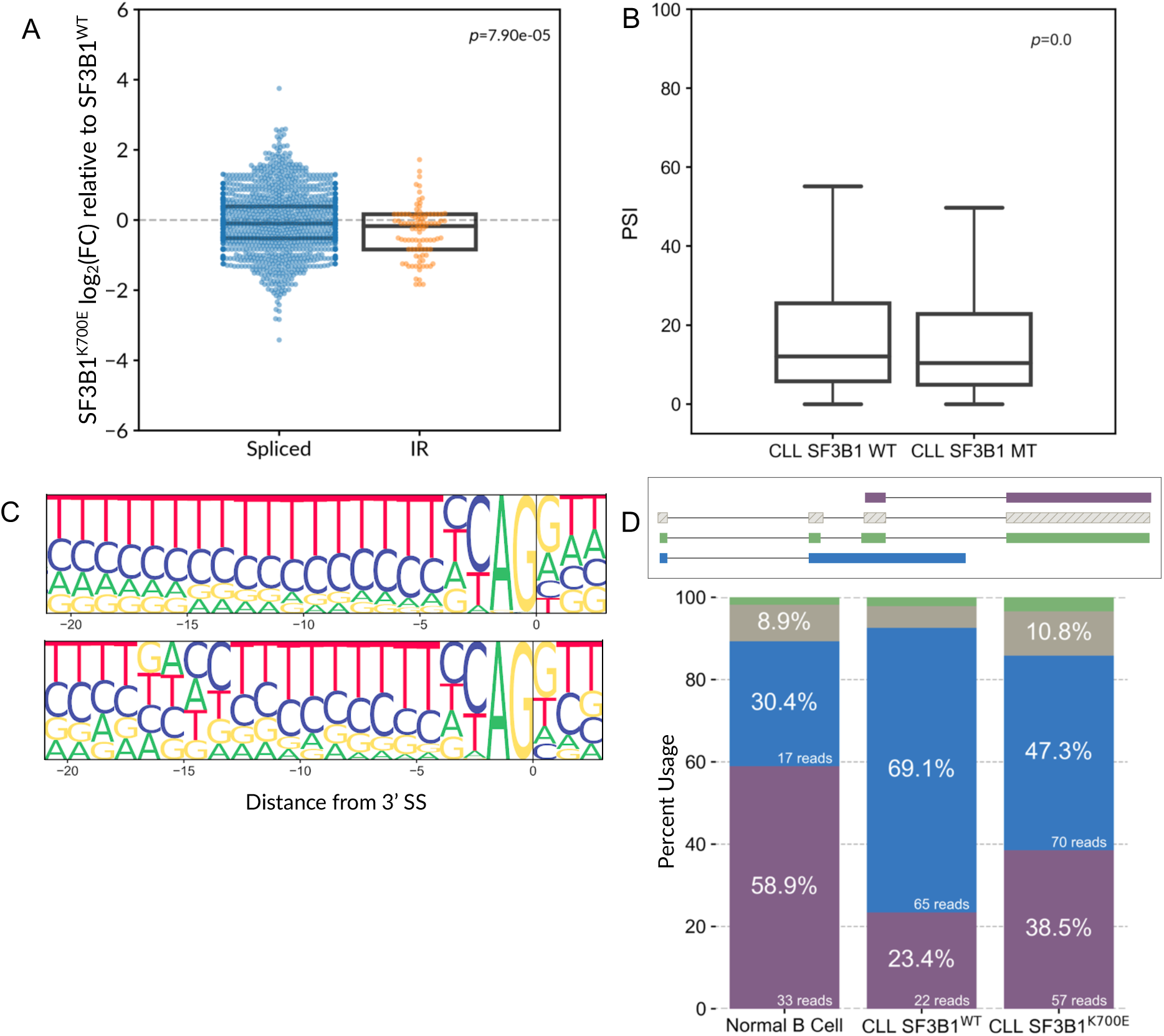
Characterization of intron retention in *SF3B1*^*K700E*^ and *SF3B1*^*WT*^. A.Expression fold-change (FC) of spliced (i.e. not intron-retaining) and intron-retaining (IR) isoforms between CLL *SF3B1*^*K700E*^ and *SF3B1*^*WT*^. B.Intron retention event PSIs reanalyzed from short-read data. C.Top: 3’ splice site motif for constitutively spliced introns. Bottom: 3’ splice site motif for significantly differentially used intron retention events between *SF3B1*^*K700E*^ and *SF3B1*^*WT*^ identified from short-read sequencing (false discovery rate < 10%). *D.DAZAP2* isoform usage. The gene structure schematic and isoform proportion colors are matched. The gene is shown 5’ on the left and 3’ on the right, displaying isoforms with more than 5 reads across all samples. Solid or hatched isoform denotes productivity or unproductivity, respectively. Raw read counts and percent usage are shown in the stacked bar graph.

We then looked at splice site motifs for alternative 3’ splice sites and differentially retained introns from short- and long-read data (Supplementary Fig. 4A-E). Using 65 alternative 3’ splice sites significantly associated with *SF3B1* mutation identified in short-read data from a cohort of 37 CLL samples (24 *SF3B1*^*WT*^ and 13 with an *SF3B1* mutation) [8], we found a tract of As 13-16 bp upstream of the canonical 3’SS, concordant with previous publications [9,10] (Supplementary Fig. 4B). Additionally, in 67 intron retention events significantly associated with *SF3B1* mutation from this same cohort we found a strong TG<u>A</u>C branch point motif [35] ∼15 bp upstream of the 3’ SS, consistent with the position of strong branch point motifs upstream of alternative 3’ splice sites that are associated with *SF3B1* mutation (Fig. 4C, Supplementary Fig. 4C). This further supports an underlying mechanism of SF3B1^K700E^ to cause splicing at a 3’SS ∼15bp downstream of a strong branchpoint [13]. The presence of a strong branch point motif also suggests that the downregulation of intron retention events in *SF3B1*^*K700E*^ are due to more efficient splicing of these introns rather than a secondary effect due to increased degradation of IR isoforms. We did not observe the same motif using alternative splicing events identified from nanopore sequencing, which may be due to fewer events identified as differentially spliced (11 events for alternative 3’ splice site, 25 events for intron retention) as well as a smaller cohort size (Supplementary Fig. 4D,E).

Using our assembled transcripts and isoform quantification approaches, we identified 122 full-length isoforms that are differentially used between CLL *SF3B1*^*K700E*^ and *SF3B1*^*WT*^ (Fisher’s exact test, p<0.05). Forty-four (36.1%) of these events involve differentially retained introns. For example, *DAZAP2* shows increased intron retention in *SF3B1*^*WT*^ relative to *SF3B1*^*K700E*^ and B-cell (Fig. 4D). This alternative isoform with a retained intron is coupled with an alternative transcription end site, which is a type of regulated coupling that is difficult to determine with short-read data. Other examples of differential isoform usage include *HLA-B*, *CDC37*, *PSMB4*, and *CD79B* (Supplementary Fig. 5). HLA-B is a protein that is part of the human leukocyte antigen complex involved in the immune system, which exhibits increased skipping of exon 6 in conjunction with an altered 3’ splice site in CLL SF3B1^K700E^, occurring in the cytoplasmic domain of the protein (Supplementary Fig. 5A). We also observe exon-skipping in *CD79B* associated with SF3B1^K700E^ (Supplementary Fig. 5D).

To further understand these isoform-level differences caused by SF3B1^K700E^, we performed a gene ontology (GO) analysis of parent genes with intron retention events more downregulated in SF3B1^K700E^ relative to SF3B1^WT^ (Fisher’s p<0.05). No GO term was enriched after multiple testing correction; however, the most enriched terms included regulation of antigenic response to inflammatory stimulus (p=2.18e-4), MAPK cascade (p=5.42e-3), signal transduction by protein phosphorylation (p=5.73e-3), and regulation of ERK1/ERK2 signaling cascade (p=1.36e-2) (Supplementary Fig. 6, Supplementary Table 1). As perturbed kinase signaling has been implicated in introns detained in the nucleus in glioblastoma [37], we decided to further assess whether these intron retention events could be considered detained introns.

### Productivity analysis of transcripts associated with SF3B1^K700E^ in CLL

With full-length cDNA sequencing, we are also able to more confidently predict transcripts with premature termination codons and estimate the proportion of unproductive transcripts. We define unproductive isoforms as those that have a premature termination codon that is 55 nucleotides or more upstream of the 3’ most splice junction (Fig. 5A) [38]. Although we define unproductive isoforms based on known rules for targets of the nonsense-mediated decay (NMD) pathway [39], these “unproductive” transcripts could be retained in the nucleus and, thus, not subject to NMD-mediated degradation. For example, *SRSF1* has a known transcript that is nuclear retained [40], and is one example of an “unproductive” transcript which we observe in our nanopore data (Supplementary Figure 7A). The majority of isoforms (68%) with intron retention events were determined to be unproductive (Fig. 5B, Supplementary Fig. 5B,5C,7B, Supplementary Table 2), suggesting that these are detained introns that are localized to the nucleus. We then analyzed changes in productivity, and associated IR, by focusing on differentially used isoforms in the *SF3B1*^*WT*^ sample versus the *SF3B1*^*K700E*^ sample (p<0.05). Although both unproductive and productive IR isoforms are downregulated in *SF3B1*^*K700E*^ (Fig. 4A), the reduction is observed more strongly in the productive IR isoforms (Fig. 5C) (Mann-Whitney U p=5.23e-5). The productive IR isoforms consist of IR occurring between coding exons or in UTRs. It is plausible that the IR events happening between coding exons produce aberrant protein despite being in-frame and labeled as productive. It is also possible that these “productive” IR isoforms are retained in the nucleus since they are not fully processed. Interestingly, we also observe down-regulation of unproductive, but fully spliced transcripts. The rules of NMD targeting have been used to associate the function of *SF3B1*^*MUT*^-associated splicing events from short-read data [8,10]; however, our productivity analysis of full-length transcripts provides caution for the interpretation of these analysis, given that we cannot resolve which transcripts are, in fact, nuclear-retained.

**Figure 5:**
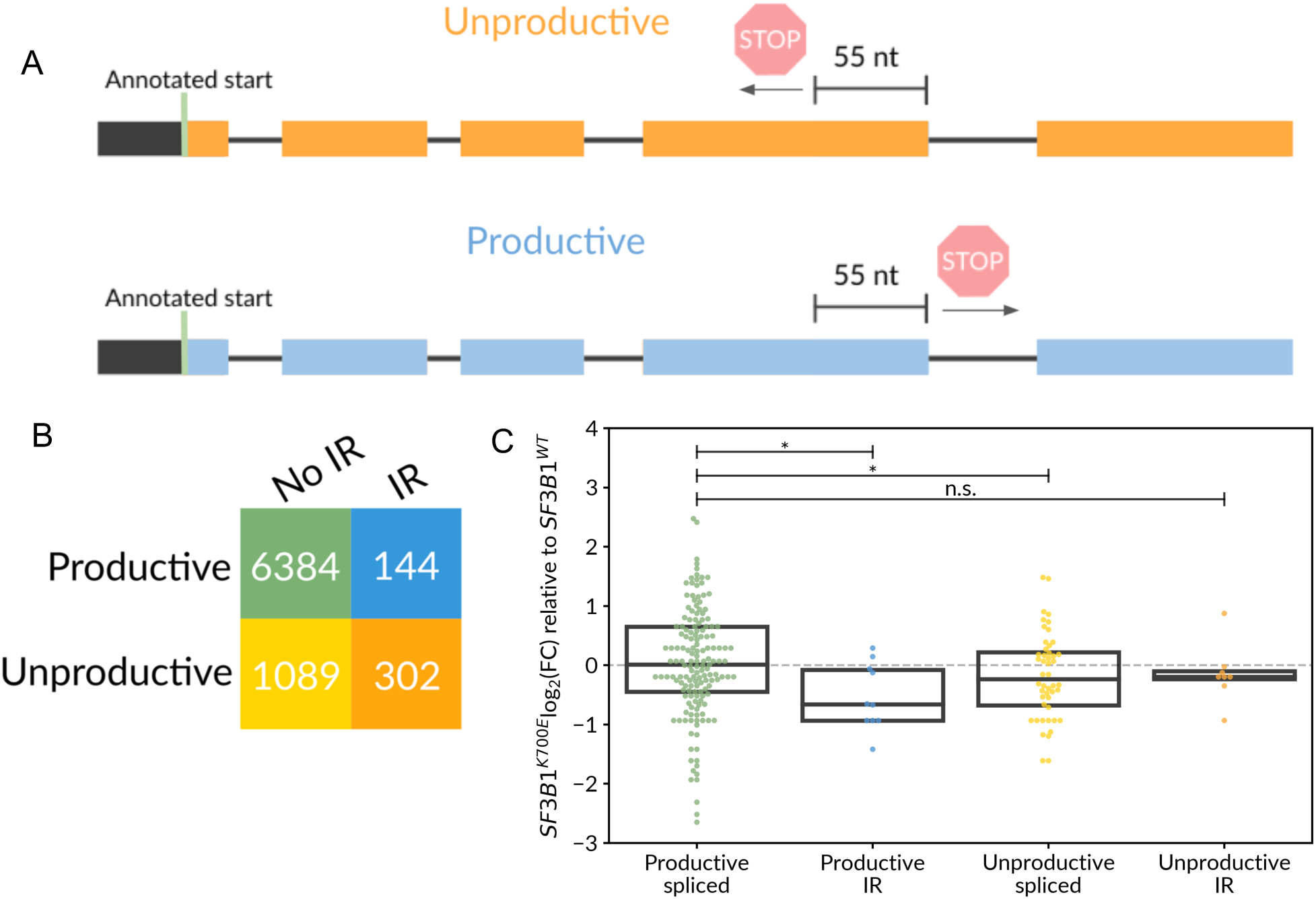
Productivity analysis of transcripts associated with *SF3B1*^*K700E*^. A.Diagram illustrating isoform productivity criterion. Isoforms from full-length reads are translated starting from annotated start codons. Isoforms with premature stop codons (PTCs) found upstream of the last splice junction are considered unproductive and downstream of the last splice junction are productive. B.Table of isoform productivity and intron retention event contingency in CLL data. C.Expression fold-change (FC) comparing 122 differentially used isoforms between CLL SF3B1^K^700^E^ to SF3B1^WT^. Isoforms of the parent genes of the differentially used isoforms are plotted according to the same color scheme as 5B. An asterisk above lines comparing categories denotes the significance of a Mann-Whitney U test, p<0.05.

## Discussion

In this study, we identified splicing changes in the context of full-length isoforms in primary CLL samples. Long-read sequencing data provided more confidence in identification of intron retention events, revealing downregulated retained introns in *SF3B1*^*K700E*^ relative to *SF3B1*^*WT*^. More generally, we observed increased intron-retaining isoforms in CLL samples that are *SF3B1*^*WT*^ when compared to a normal B-cell sample. Introns more significantly spliced out in *SF3B1* mutated samples contained a strong branch point TG<u>A</u>C sequence ∼15 bp upstream of the 3’ splice site, consistent with previously reported branch site motifs of altered 3’ splice sites associated with the mutation [13]. Gene ontology analyses revealed kinase signaling and inflammatory response genes associated with downregulated intron retention events in SF3B1^K700E^. As the majority of these IR isoforms are unproductive, we postulate that many of these are detained introns of transcripts that are kept in the nucleus rather than exported and subjected to degradation through the NMD pathway.

Using nanopore sequencing data, we were also able to identify alternative splice site usage associated with *SF3B1*^*K700E*^. These splice site changes were consistent with alterations identified using short-read data; however, the number and statistical significance of differentially spliced events identified was limited by our nanopore read depth and cohort size of this study. The difficulty of detecting subtle splicing alterations in nanopore reads can by helped by our workflow to realign reads to collapsed transcripts, for better splice site read coverage. However, variability in sequencing depth observed between flow cells affected this analysis, indicating the importance of increased sequencing depth and multiplexing to avoid batch effects. Additionally, our stringent filters in analyzing splice junctions potentially resulted in fewer significant alternative 3’ and 5’ splice site events identified in nanopore data compared to short-read sequencing. That is, we removed any novel, nanopore-only splice sites from our analysis due to limited evidence of their validity. The errors in nanopore reads pose a challenge for genomic alignment, even after optimization of the alignments for splice correctness, and many of these novel splice sites may not represent truly novel results but are artifacts of misalignment.

In what the nanopore lacks in accuracy, it compensates for in length. Our methods have been evaluated to address the low accuracy by defining confident isoforms with accurate splice sites. FLAIR adds to the relatively new space of tools for working with nanopore data and analyzing splicing differences. Although no short-read data is required for FLAIR, it can be used with matched short-read data for increased splice site accuracy. FLAIR does not require high-accuracy long reads, as it is designed to create consensus isoforms from reads with high error rates. Using nanopore long-read data, we are equipped for better quantification at the isoform-level. In addition, an improved intron retention and transcript productivity analyses that long-read data provides was important for promoting our understanding of SF3B1 biology in CLL.

The downregulation of productive IR transcripts observed in CLL *SF3B1*^*K700E*^ relative to *SF3B1*^*WT*^ both globally and within differentially used genes, in conjunction with downward shifts of unproductive spliced and unproductive IR transcripts, may represent a shift in increased relative abundance of productive spliced isoforms in *SF3B1*^*K700E*^. However, we are not able to distinguish between nuclear-retained or cytoplasmic transcripts; therefore, our productivity analysis is only a prediction. As *SF3B1*^*K700E*^ is more effective at splicing out retained introns from kinase signaling related genes (Supplementary Fig. 6, Supplementary Table 1), it is conceivable that the *SF3B1*^*K700E*^ mutation causes a more active signaling state [37].

A previous publication with short-read sequencing shows that *SF3B1* mutations cause cryptic splice site usage, predicted to produce unproductive spliced isoforms [10]. From our data, we observe a decrease in full-length spliced transcripts predicted to be unproductive. As short-read sequencing has greater depth, it is easier to detect unproductive transcripts, many of which can be lowly expressed due to NMD degradation. Our nanopore sequencing runs yielded fewer than 200,000 pass reads per sample, making it difficult to detect lowly expressed transcripts. The more highly expressed unproductive transcripts we detected with low-depth nanopore sequencing could represent nuclear retained transcripts, motivating future studies that perform cellular fractionation to test this hypothesis.

The throughput of nanopore technology has increased with subsequent iterations of flow cells and we expect depth to be less of an issue going into the future. Despite the lack of depth, our study demonstrates the ability of the nanopore to identify and quantify cancer-specific splicing and isoforms. Future cancer full-length cDNA or native RNA sequencing [41] with increased depth and a larger cohort will likely lead to additional insights into which transcripts are likely protein-coding or unproductive leading to more informative follow-up functional studies. These alternative full-length transcript variants may also function as biomarkers of potential prognostic or therapeutic relevance.

## Acknowledgements

We would like to acknowledge Ashley Byrne, Miten Jain, and Christopher Vollmers for their help with nanopore library preparation and sequencing. We would also like to thank Mark Akeson for helpful feedback. This work was supported by the Damon Runyon Cancer Research Foundation to A.N.B. A.D.T. was funded through NIH grant 5T32HG008345. A.D.T was partially supported by R01HG010053 (PI: Mark Akeson, UC Santa Cruz). C.M.S. was supported by training grants NIH T32GM008646 and 2R25GM058903.

## Author Contribution

A.D.T. and A.N.B. designed the study. A.D.T., C.M.S., and E.H. designed experiments. A.D.T. performed experiments. A.D.T., C.M.S., M.J.B., K.H., and A.N.B. wrote code and analyzed data. A.D.T., C.M.S., M.J.B., K.H., C.J.W., and A.N.B. interpreted the data. C.J.W. provided donor samples. A.D.T. and A.N.B. wrote the manuscript with input from all other co-authors.

## Methods

### Data generation and handling

RNA was extracted from tumor samples using methods described in Wang et al. [8]. The sample IDs of the CLL *SF3B1*^*WT*^ and *SF3B1*^*K700E*^ samples are DFCI-5047(CLL024) and DFCI-5067, respectively. The extracted RNA was reverse transcribed using the SmartSeq protocol [42] and cDNA was PCR-amplified, adding ISPCR adapters to the ends of all amplicon sequences as described in Byrne et al. [22]. Oxford MinION 2D amplicon libraries were generated according to the Nanopore community protocol using library preparation kit SQK-LSK208 and sequenced on R9 flowcells. Basecalling was performed with albacore v1.1.0 2D basecalling using the --flowcell FLO-MIN107 and --kit SQK-LSK208 options. 2D reads were used for consequent analyses. ISPCR adapters were removed using porechop 0.2.3 [43].

### Spliced alignment and splice site correction

Reads were aligned to hg38 using minimap2 v2.7-r654 [26] in spliced alignment mode with the command ‘minimap2 -ax splice’. After alignment, gaps of 30 nt were smoothed using mRNAtoGene [44]. After alignment, splice sites were iteratively assessed for validity. Splice sites were corrected if they were within a 10-nt window of an annotated splice site using the GENCODE v24 annotation or splice site observed in the matched short-read data. Novel splice sites identified solely through nanopore sequencing were not considered valid if the sequence motif was not the canonical GT-AG splice motif. Isoforms were splice-corrected with wiggle size of 10 to the nearest valid splice site, as determined by orthogonal short-read data, genomic sequence, and annotated junctions.

### Alternative 3’ and 5’ Splice Site Analysis

Genome-aligned nanopore reads were splice-corrected and any reads with novel splice sites were removed. The reads were then collapsed with a threshold of 1 to produce a comprehensive set of isoforms for increased sensitivity to splice site changes. Nanopore reads were realigned to this set of isoforms to obtain better isoform counts for which junction-level quantification could be extrapolated from. The weighted-identity method was used to quantify isoforms for this analysis to increase the number of reads counted while minimally introducing read-isoform assignment errors. A Fisher’s exact test was used to detect alternative 3’ and 5’ splice junction events between two samples, with the constraints that each junction possesses total coverage of at minimum 5 reads, each aberrant junction possess coverage of at minimum 3 reads, and that alternative splice sites be no more than 200nt apart.

### Data Simulation

We created a wrapper script for NanoSim to generate simulated transcriptomic data. We simulated reads from transcripts of genes that were found to be expressed in our real nanopore data, with read lengths modeled after real data using our wrapper script. We used the hg38 nanopore error model simulated from step 1 of NanoSim using nanopore 2D sequencing reads.

### Isoform Assembly

We used Mandalorion v0.2 [22] and FLAIR v1.0 to assemble isoforms using reads from all runs using the same values or equivalent parameters. For both, we used a transcription start/end site (TSS/TES) window of 20 nt and a minimum number of 3 supporting reads per isoform. For FLAIR, to assemble the first-pass assembly, TSS/TESs were determined by the density and frequency of observed sites. We compared 20 nt windows of TSSs and picked the most frequently represented site in the TSS window, with the maximum number of windows picked set at 2 and the minimum number of reads in the window set at 1. TESs were determined for each TSS window using the TESs of reads in the TSS window in the same manner, picking the most frequent TES in the TES window with the most end sites. The final nanopore-specific reference isoform assembly is made by aligning raw reads to the first-pass assembly transcript sequence using minimap2, keeping only the first-pass isoforms with a minimum number of 3 supporting reads.

### Weighted read assignment for isoform quantification

We tested three approaches to isoform quantification. Isoforms were quantified by first aligning nanopore fastq sequences to transcriptome sequences of the corresponding assembled isoform using minimap2 v2.7-r654 [26] with default settings and sam output. One approach considered only unique read alignments, with each unique read alignment being one count for its respective transcript. Another approach used only primary alignments, picking the best alignment for each read, determined by minimap2. The weighted-identity approach counts unique read alignments as a single count and distributes reads with non-unique alignments amongst each alignment. That is, reads with multiple alignments were counted by first calculating the length weighted-identity scores by multiplying (1) the identity (number of matches divided by length of aligned read) of each alignment with (2) the length of the aligned transcript. The scores were all normalized to sum to 1, with the normalized scores representing the reads’ contribution to each aligning transcript.

### NMD and Intron Retention Analysis

Fold change was calculated using counts of isoforms in both conditions normalized by library size. For identification of NMD-sensitive transcripts, we used annotated start codons and translated the full-length assembled isoforms. If a transcript overlapped more than one annotated start codon, the 5’ most start codon was selected. A premature termination codon was defined as a stop codon detected before 55 nucleotides or more upstream of the last splice junction [38]. Introns were defined as spanning gaps greater than 100 nt with confident splice junctions, and we classified retentions as having an isoform spanning the majority of another isoform’s intron within 10 nt on both ends of the intron.

### GO analysis

GO analysis was performed with the R package goseq v1.32.0 [45], setting the parameter method=hypergeometric to remove the correction for gene length bias that affects short-read data. GO terms with only one term in the category were removed from further analysis.

## Code availability

Scripts developed from this work can be found at https://github.com/BrooksLabUCSC/FLAIR. These scripts include flair.py, which can be used to run FLAIR. The individual scripts that comprise flair are sam_to_psl.py, correctSplice.py, collapse_isoforms_precise.py, and find_alt_3ss.py for sam-to-psl conversion, splice site correction, isoform collapsing, and alternative splice site testing respectively. In addition, the NanoSim_Wrapper.py script, a wrapper for simulating transcript reads using the nanopore genomic read simulator Nanosim, can be found at https://github.com/BrooksLabUCSC/labtools.

---

**Supplementary Figure 1:**
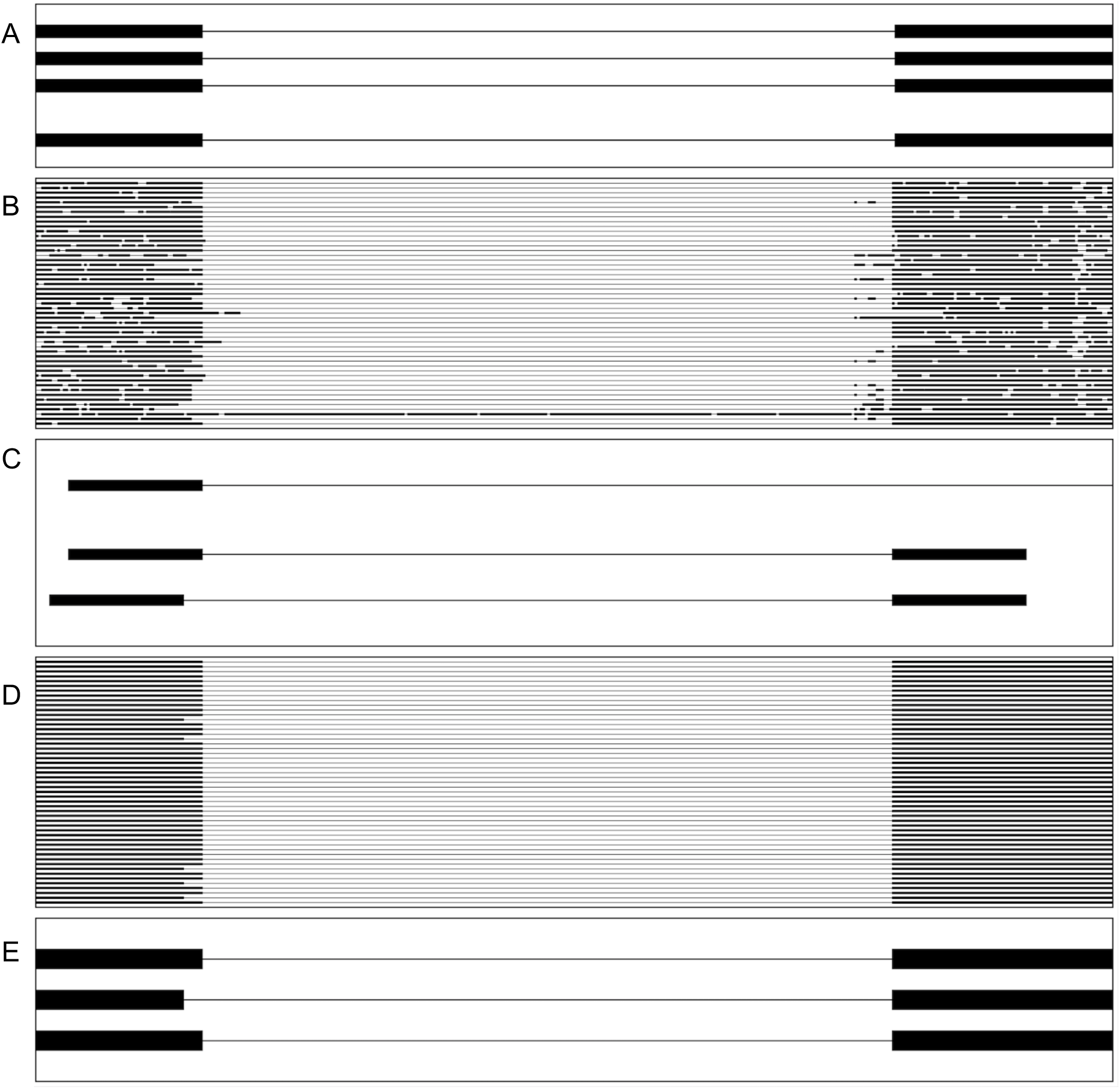
Correcting splice junctions in nanopore read alignments. A.View of a single junction for all isoforms of *FTH1* as shown in GENCODE v24 basic annotation. All following figures are of the same junction as A. B.A subset of the nanopore reads aligned to the genome using minimap2. C.Splice junctions observed in matched short read data D.Splice- and gap-corrected nanopore reads using splice junctions from A and C. E.First-pass isoform assembly by collapsing reads with the same splice junctions.

**Supplementary Figure 2:**
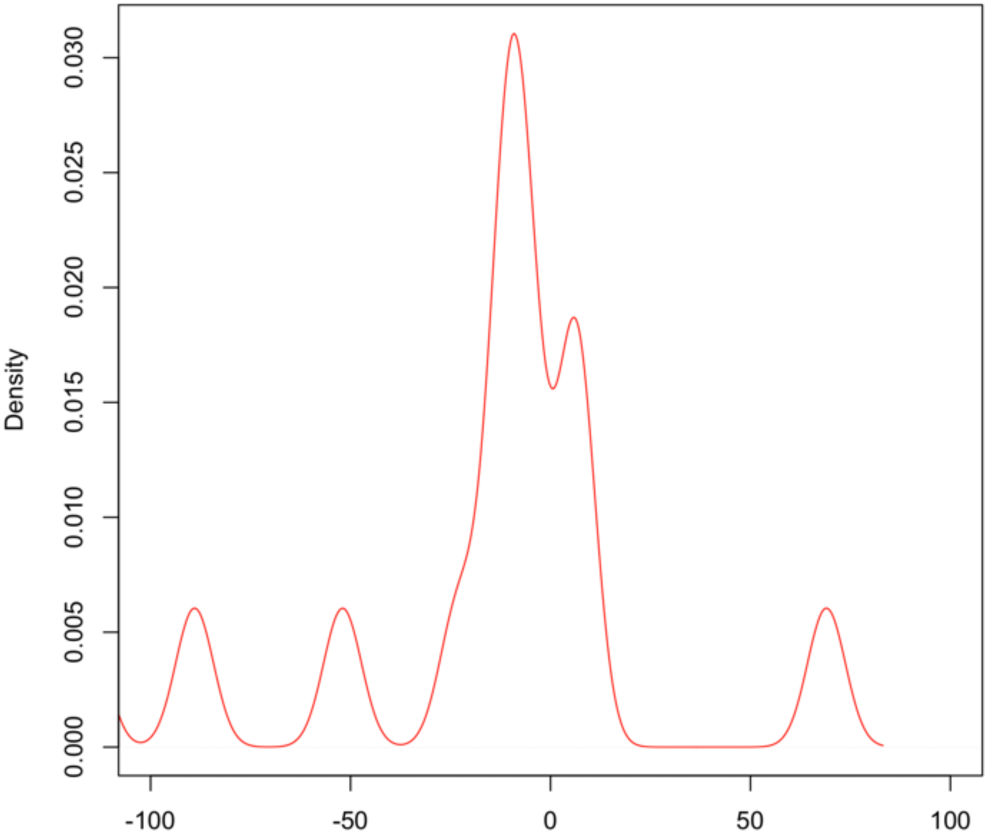
Distance of altered 3’ splice sites associated with *SF3B1*^*K700E*^. Density distribution of alternative 3’ SS used significantly more in SF3B1^K^700^E^ (n=14).

**Supplementary Figure 3:**
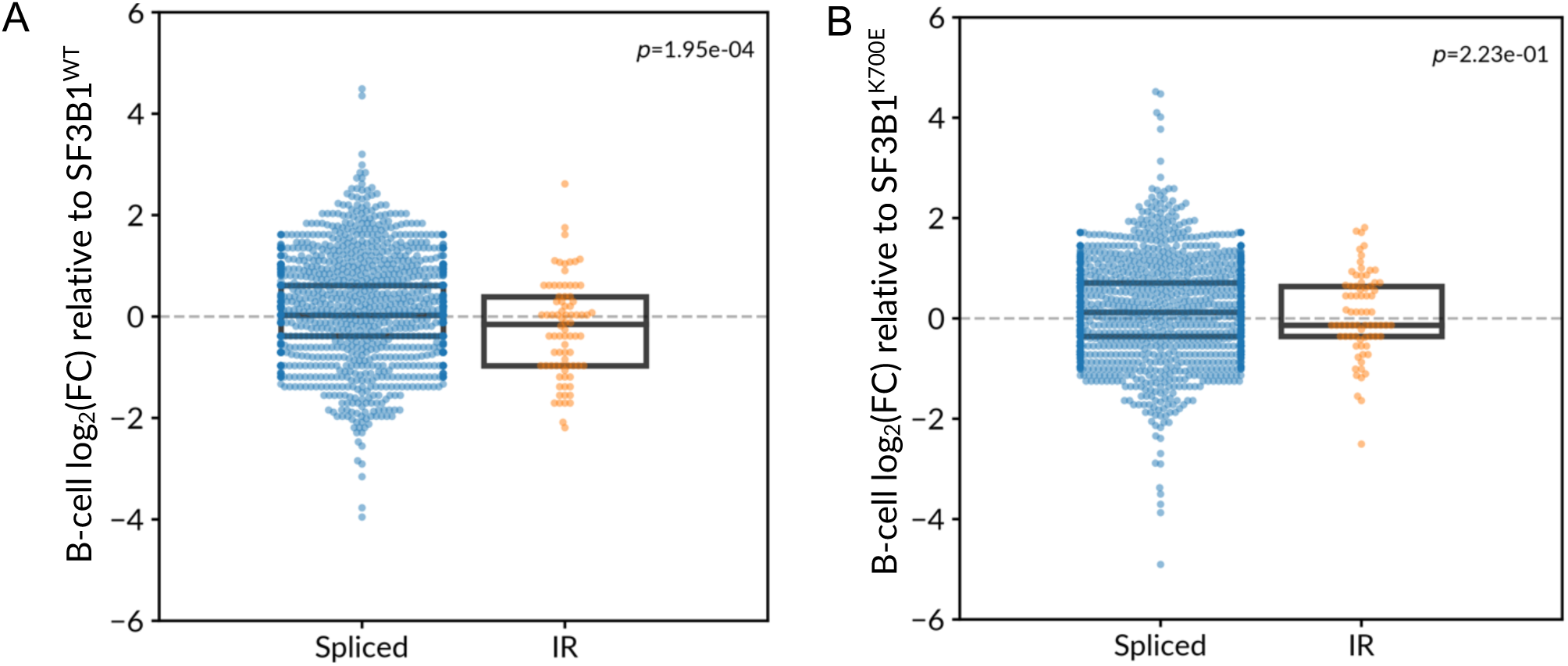
Comparison of intron retention expression. A.Expression fold-change of isoforms between CLL *SF3B1*^*WT*^ and normal B-cell categorized as spliced (non-intron-retaining) and IR (intron-retaining). P-values shown are from a Wilcoxon rank-sum test. B.Same as A but between CLL *SF3B1*^*K700E*^ and normal B-cell.

**Supplementary Figure 4:**
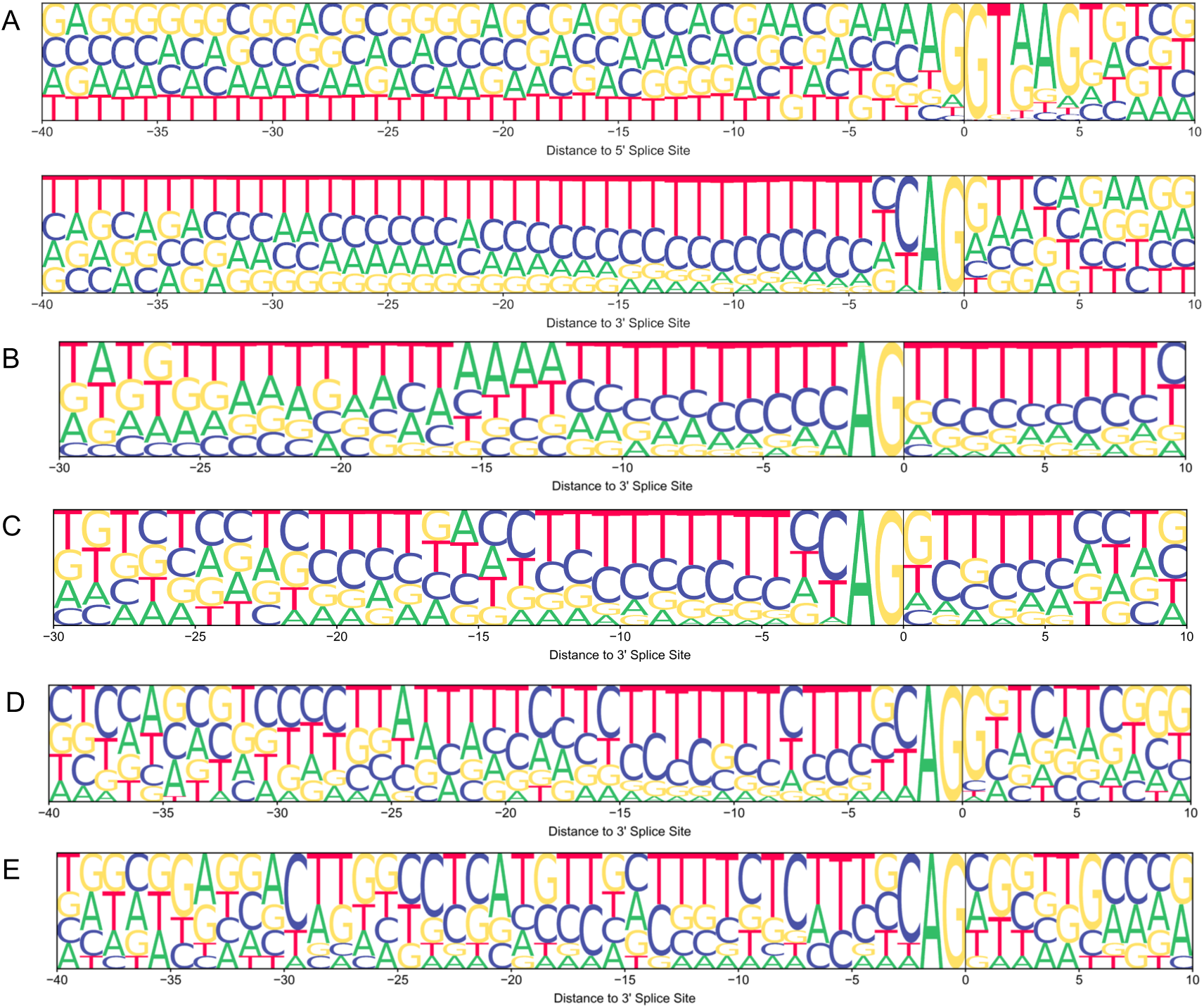
Splice site motifs of differentially splice events associated with *SF3B1* mutation. A.Splice site motifs for spliced junctions in intron-retaining isoforms identified from nanopore. Top: 5’ SS; bottom: 3’ SS. B.3’ SS motif using short-read data for significantly altered 3’ SS between *SF3B1*^*K700E*^ and *SF3B1*^*WT*^ C.Same as B using short-read data showing the 3’SS motif for IR events significantly altered (PSI *SF3B1*^*WT*^ > PSI *SF3B1*^*K700E*^) D.Same as C but using nanopore intron-retaining junctions that are more expressed in *SF3B1*^*WT*^ E.Same as B but using nanopore-determined sites that are altered 3’ SS between *SF3B1*^*K700E*^ and *SF3B1*^*WT*^

**Supplementary Figure 5:**
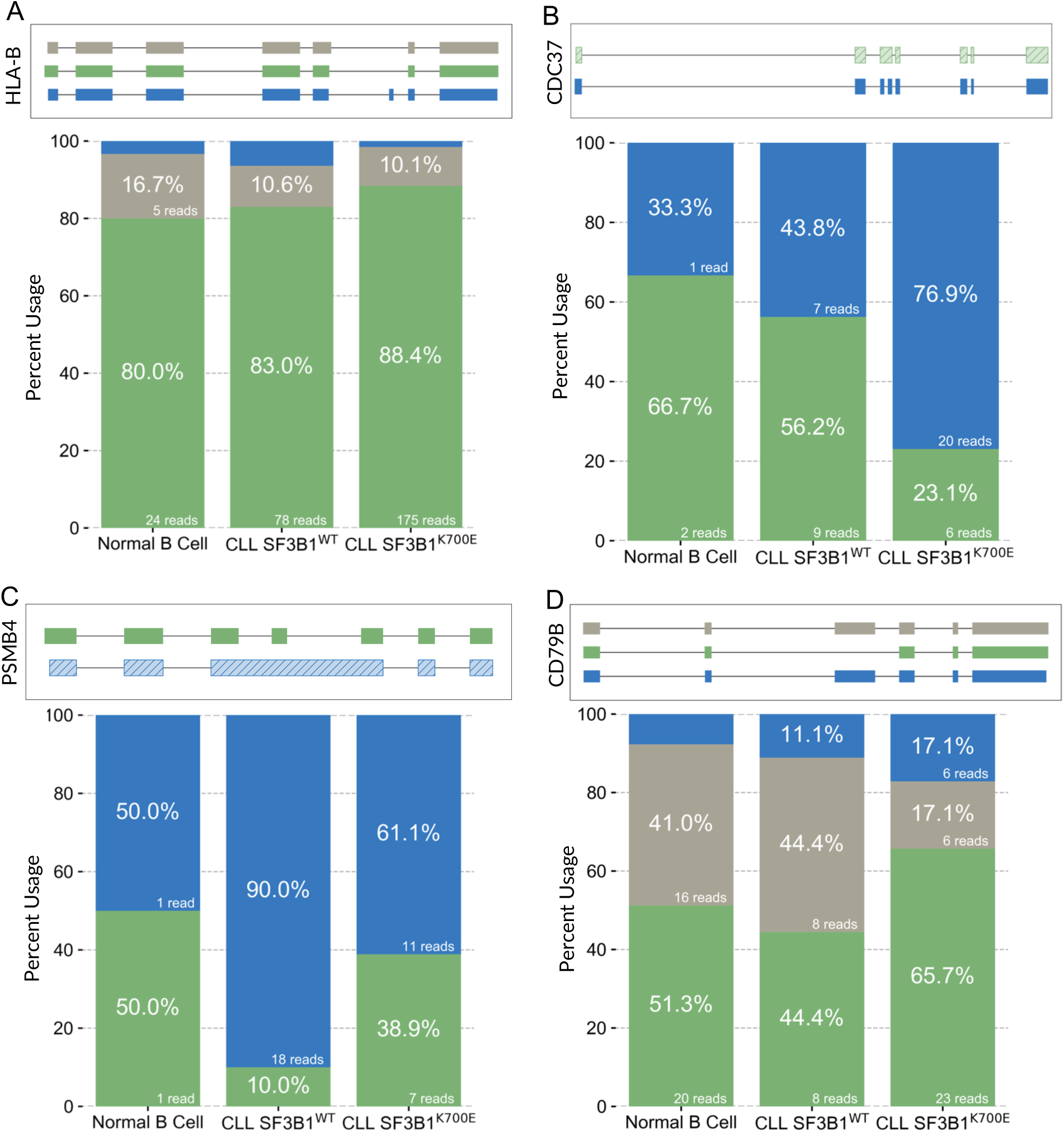
Differential isoform usage between *SF3B1*^*K700E*^ and *SF3B1*^*WT*^. FLAIR isoforms for (A) *HLA-B*, (B) *CDC37*, (C) *PSMB4*, and (D) *CD79B*. As in the main text, hatched isoforms denote unproductivity, raw read counts and percent usage are reported on appropriately sized bars, and genes are plotted 5’->3’ from left to right.

**Supplementary Figure 6:**
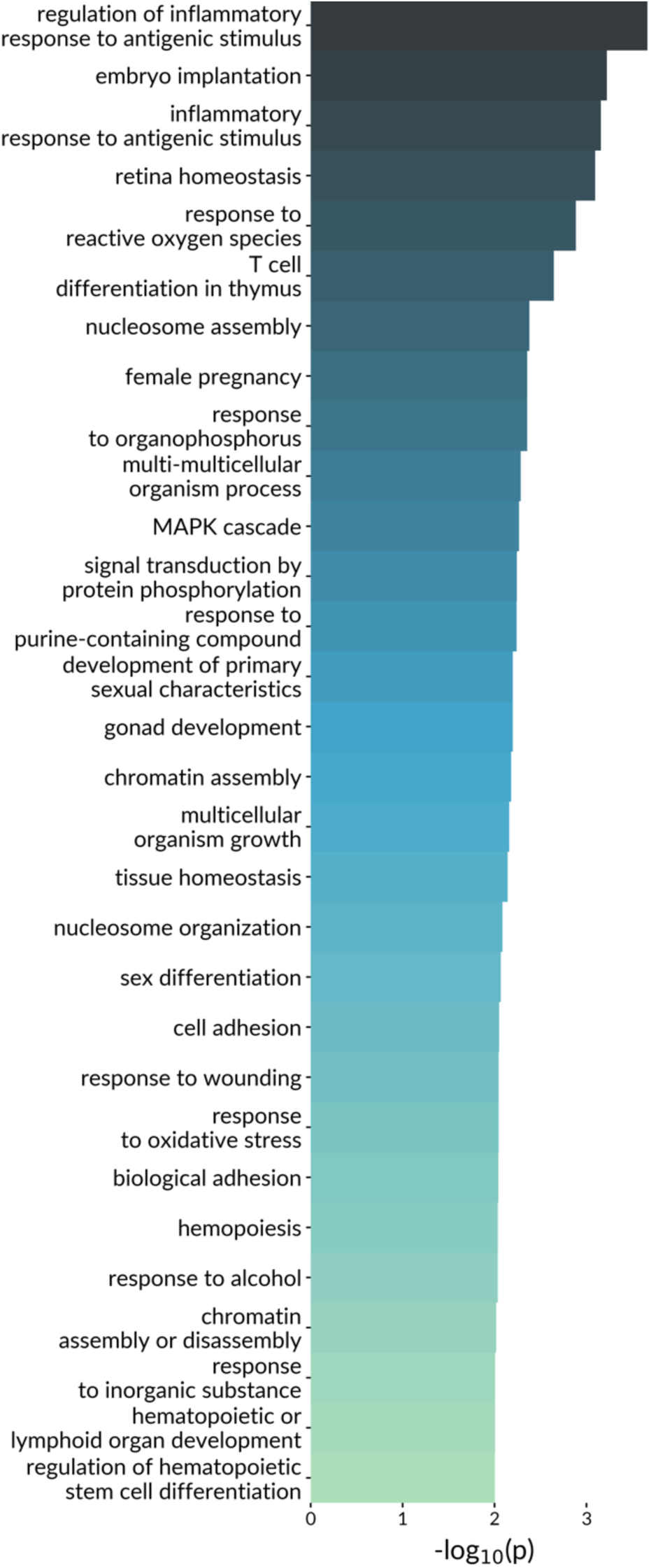
GO term analysis. Top 30 most significant (p<0.01) GO terms resulting from genes with intron retentions downregulated in *SF3B1*^*K700E*^ compared to *SF3B1*^*WT*^ (Fisher’s p<0.05). P-values shown are overrepresented categories calculated from goseq. **Supplementary Figure 7**

**Supplementary Figure 7:**
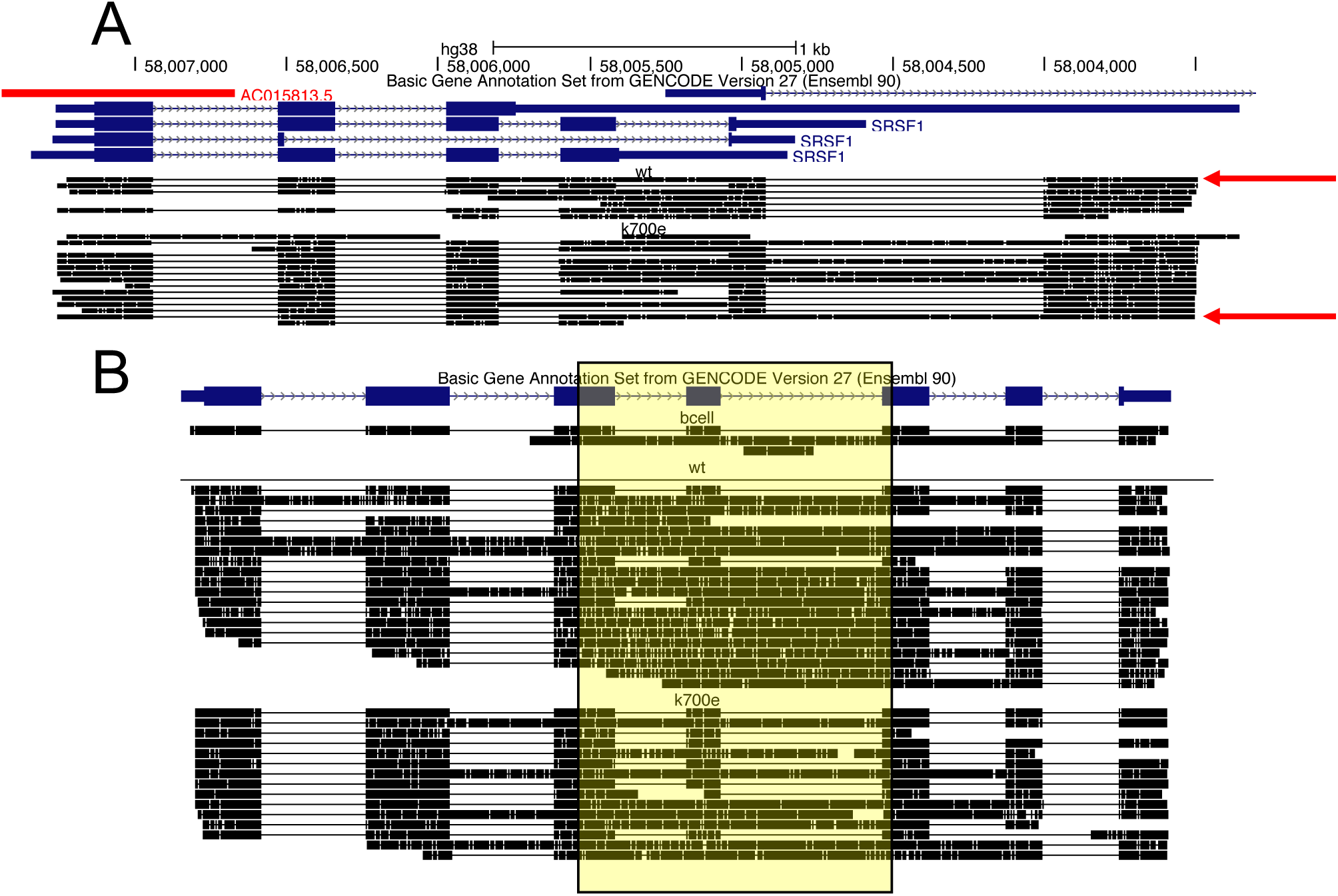
Nuclear-retained or unproductive transcripts supported by nanopore reads. A.Genome browser shot displaying annotated *SRSF1* isoforms (blue) and nanopore reads in the the *SF3B1*^*K700E*^ and *SF3B1*^*WT*^ samples (black), with the 5’ end of the transcript on the left. No reads were sequenced in B-cell. for this gene. Example reads from known nuclear-retained isoforms are indicated (red arrows). B.Same as A for *PSMB4*.

**Supplementary File 1: FLAIR transcript models derived from nanopore sequencing data.** GTF file containing isoforms assembled from the FLAIR pipeline using all nanopore data generated from this study with a supporting read threshold of 3.

**Supplementary Table 1: GO enrichment of genes with intron retention downregulated in *SF3B1***^***K700E***^ **relative to *SF3B1***^***WT***^ **determined by nanopore data.** All GO terms with p < 0.05, excluding GO terms with only 1 gene.

**Supplementary Table 2: Isoform annotations in an extended PSL format for isoforms assembled from the FLAIR pipeline.** Additional columns after column 21 containing information pertinent to this study. Spliced or intron retaining-isoforms are indicated with a 0 or 1 in column 22; productive, unproductive, and lncRNAs are marked with 0, 1, and 2 in column 23; the CLL *SF3B1*^*K700E*^, CLL *SF3B1*^*WT*^, and B-Cell samples isoform expression levels are in columns 24, 25, and 26.

